# QuASAR-MPRA: Accurate allele-specific analysis for massively parallel reporter assays

**DOI:** 10.1101/105627

**Authors:** Cynthia A. Kalita, Gregory A. Moyerbrailean, Christopher Brown, Xiaoquan Wen, Francesca Luca, Roger Pique-Regi

**Affiliations:** Center for Molecular Medicine and Genetics, Wayne State University, Detroit, MI; Department of Genetics, University of Pennsylvania, Philadelphia, PA; Department of Biostatistics, University of Michigan, Ann Arbor, MI; Department of Obstetrics and Gynecology, Wayne State University, Detroit, MI

## Abstract

**Motivation:** The majority of the human genome is composed of non-coding regions containing regulatory elements such as enhancers, which are crucial for controlling gene expression. Many variants associated with complex traits are in these regions, and may disrupt gene regulatory sequences. Consequently, it is important to not only identify true enhancers but also to test if a variant within an enhancer affects gene regulation. Recently, allele-specific analysis in high-throughput reporter assays, such as massively parallel reporter assays (MPRA), have been used to functionally validate non-coding variants. However, we are still missing high-quality and robust data analysis tools for these datasets.

**Results:** We have further developed our method for allele-specific analysis QuASAR (quantitative allele-specific analysis of reads) to analyze allele-specific signals in barcoded read counts data from MPRA. Using this approach, we can take into account the uncertainty on the original plasmid proportions, over-dispersion, and sequencing errors. The provided allelic skew estimate and its standard error also simplifies meta-analysis of replicate experiments. Additionally, we show that a beta-binomial distribution better models the variability present in the allelic imbalance of these synthetic reporters and results in a test that is statistically well calibrated under the null. Applying this approach to the MPRA data by Tewhey *et al.* (2016), we found 602 SNPs with significant (FDR 10%) allele-specific regulatory function in LCLs. We also show that we can combine MPRA with QuASAR estimates to validate existing experimental and computational annotations of regulatory variants. Our study shows that with appropriate data analysis tools, we can improve the power to detect allelic effects in high throughput reporter assays.

**Availability:** http://github.com/piquelab/QuASAR/tree/master/mpra

## 1 INTRODUCTION

Genetic variants in non-coding regions are responsible for inter-individual differences in molecular and complex phenotypes. Quantitative trait loci (QTLs) for molecular and cellular phenotypes (Dermitzakis, 2012) have been crucial in providing stronger evidence and a better understanding of how genetic variants in regulatory sequences can affect gene expression levels (Stranger, 2007; Gibbs *et al.*, 2010; Melzer *et al.*, 2008; Cheung *et al.*, 2003; Brem *et al.*, 2002). However, eQTL studies have severe limitations in identifying the true causal variant, due to linkage disequilibrium (LD) limiting the resolution of analysis. The availability of extensive functional annotations (Consortium, 2012; Pique-Regi *et al.*, 2011; Hoffman *et al.*, 2012; Moyerbrailean *et al.*, 2016b) enables the integration of functional genomic information into eQTL analysis, which can be useful to dissect the causal variant and the functional basis of the observed associations (Gaffney *et al.*, 2012; Veyrieras *et al.*, 2008; Lee *et al.*, 2009; Lappalainen *et al.*, 2013; Kichaev *et al.*, 2014; Wen *et al.*, 2015; Pickrell, 2014). SNPs that fall within a transcription factor (TF) binding site (TFBS) represent a major mechanism underlying eQTLs (Degner *et al.*, 2012). Recently, additional computational and experimental techniques have been developed to predict and detect allelic effects of SNPs in TFBS using DNase I footprinting and ChIP-seq data (from the ENCODE and Roadmap Epigenome projects) (Moyerbrailean *et al.*, 2016b; Lee *et al.*, 2015; Maurano *et al.*, 2015; Zhou and Troyanskaya, 2015). Still, it is a challenge to further validate if allelic effects in binding translate to effects on gene transcription. While all these existing computational annotations are useful for predicting the causal SNP in an eQTL, they do not prove the SNP is truly causal, nor do they properly quantify its effect on gene expression.

To dissect regulatory sequences and compare genetic effects on gene expression, different versions of high throughput reporter assays have emerged in the recent years. These include massively parallel reporter assays (MPRA) Melnikov *et al.* (2012); Kwasnieski *et al.* (2012) and self transcribing active regulatory regions sequencing (STARR-seq) Arnold *et al.* (2013) that can simultaneously measure the regulatory function of thousands of constructs at once. MPRAs utilize a multitude of unique synthesized DNA oligos that are associated with barcodes, cloned in a reporter plasmid and transfected into cells. The transcripts are then isolated for RNA-seq. The number of barcode reads in the RNA over the number of barcode reads from the plasmid DNA is used as a quantitative measure of expression driven by the synthetic enhancer region (Melnikov *et al.*, 2012; Kwasnieski *et al.*, 2012; Patwardhan *et al.*, 2012; Sharon *et al.*, 2012; Kwasnieski *et al.*, 2014). MPRA and STARR-seq were originally developed to identify and validate regulatory regions, but they can also be used to compare allelic effects of genetic polymorphisms or methylation (Lea *et al.*, 2017). Recent studies used this technique to compare allelic variants of SNPs with the aim to dissect, at a large scale, the causal nucleotide in eQTL and Genome Wide Association Study (GWAS) signals. Specifically, (Vockley *et al.*, 2015) used a STARR-seq derived method (POP-STARR-seq) to measure allelic effects on gene expression for population based variation in 104 regulatory regions, and a more recent study by (Tewhey *et al.*, 2016) adapted MPRA to fine-map variants associated with gene expression in lymphoblastoid cell lines (LCLs) and HepG2.

The application of MPRA to quantify the allelic effects of regulatory variants is very similar to the challenge posed by allele-specific expression (ASE) in RNA-seq data. However, one key difference is that the proportion of plasmids for each allelic construct may not be in a 1:1 ratio. Few off-the-shelf statistical methods have been used for processing and analyzing these large MPRA datasets (Table 1), but they do not consider several technical issues that can lead to false positives, such as base-calling error and over-dispersion. As demonstrated in RNA-seq ASE approaches, a binomial distribution fails to account for overdispersion and results in overly optimistic *p*-values, while a beta-binomial distribution is a more adequate choice (Kumasaka *et al.*, 2015a; van de Geijn *et al.*, 2015). Compared to RNA-seq ASE methods that combine all reads across haplotypes, in MPRA we do not need to accommodate for the uncertainty in phasing or haplotyping as the complete sequence of each construct is known. By design we can also avoid oligonucleotides that could lead to ambiguous mapping. This is in stark contrast to using the entire human genome/transcriptome, which typically requires extensive pre-processing. This is because many genomes contain large number of repetitive and quasi-repetitive regions that are only one SNP or base-calling error away from many other paralogous regions. Here we further extend QuASAR (Harvey *et al.*, 2014), an approach which considers both over-dispersion and base-calling errors, to test for allelic imbalance in MPRA constructs when the default proportions are not equal. The new method allows for estimates of the dispersion parameter depending on variant-specific read coverage, and produces summary statistics that are easy to incorporate in downstream analyses.

**Table 1.** Statistical methods for ASE and MPRA analysis.

| Conditions | Type of test |  |  |  |  |
| --- | --- | --- | --- | --- | --- |
| | T | Fisher | Bin | β-bin | QuASAR |
| Previously used in MPRA | X | X |  |  |  |
| Previously used for ASE |  |  | X | X | X |
| Requires normally distributed data | X |  |  |  |  |
| Underestimates the effect of biological variability | X | X | X |  |  |
| Handles overdispersion |  |  |  | X | X |
| Accounts for base calling error |  |  |  |  | X |

Here we tested our new method on MPRA data from Tewhey *et al.* and we further confirmed the robustness of our method on another dataset from Ulirsch *et al.*. First we compared our new QuASAR-MPRA statistical test to other tests employed in MPRA and ASE analyses (Table 1). We then demonstrate that the QuASAR-MPRA test better calibrates the *p*-values under the null hypothesis, without sacrificing statistical power. Finally, we used the allelic effects identified by QuASAR-MPRA to investigate whether the genetic variants that fall within genomic annotations, such as TF binding motifs, are good predictors for allele-specific regulatory function. Our study shows the potential value of using robust allele-specific analysis in high throughput reporter assays, to improve fine mapping analysis of association signals and validate genomic annotations of regulatory variants.

## 2 METHODS

### 2.1 Data source and pre-processing

We downloaded processed read counts from GEO (GSE75661) ftp://ftp.ncbi.nlm.nih.gov/geo/series/GSE75nnn/GSE75661/suppl/GSE75661_79k_collapsed_counts.txt.gz (Tewhey *et al.*, 2016).This MPRA study was designed to look at ASE in 39,479 oligo pairs representing 3,642 eQTLs from the GEUVADIS RNA-seq dataset of lymphoblastoid cell lines (LCLs) from European and African individuals (Lappalainen *et al.*, 2013). It has a large number of experimental replicates (8 LCL replicates), and makes use of barcodes (an average of 73 unique barcodes per oligo per replicate) to remove PCR duplicates, making this an ideal dataset to work with. We considered separately sequences in the forward and reverse strand direction in the library, as direction of the regulatory region could potentially affect reporter gene and therefore barcode expression. Tewhey *et al.* found that filtering the data to remove variants with low coverage greatly reduced the variability between replicates. Higher variance could then lead to falsely identifying ASE. We therefore began processing the dataset by applying a counts filter. For each direction we removed all cases with less than five reads on the reference and alternate allele, and where the sum of two alleles was ≤ 100. This gave us a total of 33,664 SNPs in the DNA library as input to the RNA library.

For the RNA library, we first separated the library into forward and reverse directions, and then required that RNA constructs were in the DNA library. We used a counts filter of 5 for both reference and alternate alleles so that we were only looking at variants that had sufficient reads covering both alleles to test for allele-specific effects on expression. This left us with 19,287 SNPs in the forward library and 19,748 SNPs in the reverse library or 33,653 SNPs total represented.

We additionally applied the QuASAR-MPRA method to a separate dataset by Ulirsch *et al.*. We downloaded processed read counts from http://www.bloodgenes.org/RBC_MPRA/ (Ulirsch *et al.*, 2016). This dataset comprised of 2,756 variants in strong linkage disequilibrium with 75 sentinel variants associated with red blood cell traits, with reference and alternate alleles represented in the pool of constructs. Each variant has 3 sliding windows of coverage, which we treated as separate constructs (rather than combining counts per variant). This dataset comprised of 2 DNA and 6 RNA replicates (from K562 cells). The data was processed using the same steps as with the Tewhey *et al.* data, resulting in 2,669 SNPs in total.

### 2.2 Baseline statistical methods for comparison

To test for ASE there are several different methods available (Table 1). The *t*-test, Fisher’s exact test and binomial test are classical tests remarkably appealing due to their simplicity. However, they have several limitations, as they cannot be tuned to the context of the experiment, such as levels of overdispersion (eg. from biological and technical variability) which are known to exist in ASE data (Castel *et al.*, 2015; Skelly *et al.*, 2011; Anders *et al.*, 2010). A paired Student’s *t*-test for ASE can be used to test whether the mean expression of the reference allele is equal to the mean expression of the alternate allele. This test requires multiple replicates in order to calculate a mean for each allelic expression group that has little variance, otherwise the test will not have the power to detect differences. Fisher’s exact test has been used previously to identify ASE (Romanel *et al.*, 2015), by testing whether the reference and alternate allele counts’ proportions are the same. Rejection of the null hypothesis, however, only informs us that the difference between the average counts in the two samples is larger than one would expect between technical replicates. In the binomial test, the null hypothesis is that observed values for two categories do not deviate from the theoretically expected distribution of observations. In ASE, the binomial test is used to determine whether the ratio of the two alleles is significantly different from the expected proportion (e.g. 0.5).This is the classic test that has been employed previously to detect ASE in RNA-seq studies, and assumes that read counts within each gene are binomially distributed (Kilpinen *et al.*, 2013; Consortium *et al.*, 2015; Lappalainen *et al.*, 2013; Buil *et al.*, 2014). Even accounting for reference mapping bias in RNA-seq reads, *p*-values have been found to remain inflated (Castel *et al.*, 2015). Other methods handling ASE such as WASP, RASQUAL, EAGLE(Kumasaka *et al.*, 2015b; van de Geijn *et al.*, 2014; Knowles *et al.*, 2017) use a per SNP overdispersion parameter and give well calibrated *p*-values. However these methods perform ASE QTL mapping and their application to MPRA would require a large number of replicates (*>* 15).

To reproduce the Student’s *t*-test performed by Tewhey *et al.*, we calculated the log_2_ ratio for the reference and alternate allele constructs (RNA/DNA) for each replicate. These values were used as input for a paired *t*-test in R. To perform the Fisher’s exact test on the MPRA counts data, we first added a pseudocount of 1 (Vockley *et al.*, 2015) to each RNA and DNA reference and alternate allele counts and then used the fisher.test function in R. To perform the binomial test on the MPRA counts data, we compared the reference and alternate allele counts to the DNA proportion (reference allele/ reference allele + alternate allele). To combine the *p*-values for the two LCL individuals, we used Fisher’s method (Tewhey *et al.*, 2016).

### 2.3 QuASAR Approach

QuASAR by default assumes that under the null hypothesis of no allelic imbalance the reference and alternate allele read counts should be at 1:1 ratio. However, in MPRA, the proportion *r*_*l*_ of the reference reads is not necessarily 0.5 across all the ℓ. genetic variants, due to differences in PCR amplification, as well as cloning and transformation efficiencies. Here, we have extended QuASAR to test for differences between the proportion of reference reads in DNA *r*_*1*_ and the proportion obtained from RNA reads ρℓ. To reject the null hypothesis ρℓ. = *r*_*l*_, we extend QuASAR’s beta-binomial model. The observed reference *R*_*l*_ and alternate *A*_*l*_ allele read counts at a given ℓ. are modeled as:

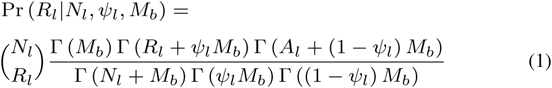

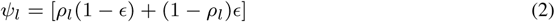

where *N*_*l*_ = *R*_*l*_ +*A*_*l*_ is the total read count at *l*, and *M*_*b*_ is the concentration parameter that controls over-dispersion of the mean proportion centered around *ψ*_*ℓ*_,which also incorporates in the model a base-calling error *ε* and the allelic ratio ρℓ. overall-mean. We can estimate *ε* using an EM procedure (Harvey *et al.*, 2014), but here for MPRA we fixed 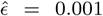 as a conservative estimate of the true base-calling error rate.

We have found previously for ASE that overdispersion decreases with greater depth of coverage (Figure S9 in Moyerbrailean *et al.* (2016a)). Therefore here, as compared to our previous implementation of QuASAR, we use different *M*_*b*_ parameters depending on the sequencing depth *N*_*l*_. We bin *N*_*l*_ into different quantiles (here deciles) and we estimate *M*_*b*_ for each bin separately using a grid search:

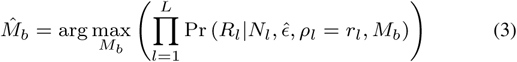

This should work well when the number of sites (i.e., SNPs tested) is relatively large so each bin *b* has *>* 200 observations to estimate *M*_*b*_. In our experience sequencing depth is a major determinant for M, and because we estimate M under the null, we tend to be conservative (i.e., M is the worst case scenario for all the constructs that belong to the same group). As a consequence, the QuASAR-MPRA *p*-values remain well calibrated (or in the worst case scenario they will tend to be slightly conservative).

We estimate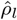 using (1) with 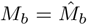 from (3) and a standard gradient method (L-BFGS-B) to maximize the log-likelihood function:

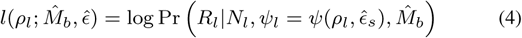

Finally, all parameters are used to calculate the LRT statistic, contrasting 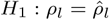to *H*_0_ ρ_*l*_= *r*_*l*_ and the resulting *p*-value.

For comparison, we performed the original QuASAR analysis on the data as well, as described in Harvey *et al.*.

### 2.4 QuASAR meta-analysis

Using the QuASAR approach, we can generate summary statistics of the allelic imbalance that can be used for downstream analyses. For example, to compare DNA to RNA, or between RNA of different cell-types, or to perform meta-analysis of multiple MPRA libraries. Instead of using an estimate of the allelic proportion ρ_*l*_, in the QuASAR approach we report the estimate of *β*_l_ = log(*ρ_l_*/(1 − *ρ_l_*)) and its standard error 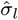 using the second derivative (i.e. Hessian) of the log-likelihood function in (4). We prefer the logistic transformed parameter *β_l_* as it provides a more robust fit and the second derivative is better behaved than that of *ρ_l_* on the edges.

To illustrate this for the Tewhey et al. data, we combined the summary statistics for the two LCL individuals using standard fixed effects metaanalysis. The effect size *β_l,n_* of each replicate n is weighted by 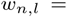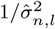, to calculate the overall effect size and standard error:

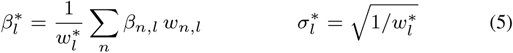

where 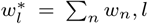We can then calculate the Z-score and *p*-value to test for an overall change between all the RNA replicates combined with respect to the original DNA proportion *β_0_*:

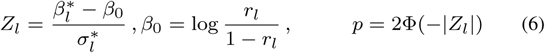

Across all the paper, *p*-values were corrected for multiple testing using the Benjamini-Hochberg’s (BH) method (Benjamini and Hochberg, 1995). To compare the different approaches we quantified the genomic inflation parameter, λ, for a set of *p*-values (Yang et al., 2011). For this we calculated the ratio of the median of the *p*-value distribution to the expected median, thus quantifying the extent of the bulk inflation and the excess false positive rate.We also use a rank sum paired test to assess statistical significance in the p-value inflation between QuASAR-MPRA and other methods with similar performance.

### 2.5 Simulations

To simulate MPRA data we randomly sampled from a beta-binomial distribution with parameters set to approximate the real data in Tewhey *et al.*. The advantage of a simulation is that we have full knowledge of which SNPs are truly imbalanced and we can empirically calculate statistical power (i.e, sensitivity) and FDR under specific assumptions. The true underlying distribution may not exactly be beta-binomial but simulations are still very useful to know how the test performs and compares to other tests. We started by simulating the DNA reads and proportions for each SNP using a beta-binomial. For this we used the DNA proportion from the Tewhey *et al.* data as the expected proportion *ρ_*l*_* and we set the concentration parameter *M* to be 200, and the total number of DNA reads *N* = 10,000.

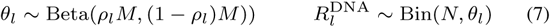

After we simulated the DNA counts *R*^DNA^, we recalculated the new DNA proportions 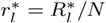. The exact value of the parameters used to generate the DNA counts are not very important and should have no effect for the simulation as the differences are captured on the RNA data once the DNA proportion is specified. To simulate the RNA data we need to simulate two conditions: 1) SNPs without ASE and the same proportion as in DNA, and 2) SNPs with ASE and a different allelic proportion than those in DNA. To do this, we explored different parameter settings for the concentration parameter *M* (10, 60 and 100), effect size Δ*β*(0.5, 1.0 and 2.0) and number of RNA replicates (2, 5 and 8). The number of reads *N*_*l*_ observed for each SNP was set up to match the average NA12878 RNA counts for each SNP (so multiple coverages are being simulated) and we divided these by a constant factor to simulate sequencing depths (1/2,1/5 and 1/10) lower than those obtained by Tewhey *et al.*. Each RNA replicate was simulated with the same parameter setting.

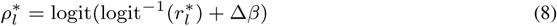

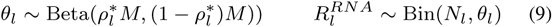

To sample from Beta and Bin, we used the `rbeta` and `rbinom` in `R` respectively. The proportion of SNPs with RNA counts with Δ*β* ≠ 0and simulated to have an allelic imbalance is 0.1% of the total SNPs. For each simulated data set we then ran the *t*-test and QuASAR and adjusted the *p*-values for multiple testing using the same BH procedure as in the real data. For each FDR control threshold we empirically calculated power (Sensitivity) and false discovery rate (eFDR). To ensure that we get robust sensitivity and FDR estimates we repeated the entire procedure 20 times and reported the average.

### 2.6 Annotation Overlap

Table S1 reports the annotations we have considered with their sources. More specifically, we considered two major sets of annotations: experimentally and computationally derived. The experimental annotations include allele-specific hypersensitivity (ASH) from (Moyerbrailean *et al.*, 2016b), dsQTLs (Degner *et al.*, 2012), and GTEx eQTLs (Consortium *et al.*, 2015).

In terms of computational annotations, a variety of different methods have been used recently to predict the allelic effect of SNP on TF binding and chromatin accessibility. GKM-svm (Lee *et al.*, 2015) uses gapped k-mer frequencies to predict the activity of larger functional genomic sequence elements, including the impact of a variant on DNase I sensitivity. It utilizes support vector machinery based on the structural risk minimization principle from statistical learning theory and kernel function which calculates the similarity between any two sequences. CATO (Maurano *et al.*, 2015) quantifies the effect of SNPs on the energy of TF binding, through overlapping SNP DHS profiles with TF motifs and applying a logistic model which takes into account site dependent features and phylogenetic conservation. DeepSEA (Zhou and Troyanskaya, 2015) uses TF binding, DHS, and histone-mark profiles with genomic sequence information as input for training a deep learning-based algorithm and predict the effects that sequence alterations have on the chromatin. DeepSEA has three major features: integrating sequence information from a wide sequence context, learning sequence code at multiple spatial scales with a hierarchical architecture, and multitask joint learning of diverse chromatin factors sharing predictive features. Finally we also used CentiSNPs, an annotation that we recently developed (Moyerbrailean *et al.*, 2016b) that uses the CENTIPEDE framework (Pique-Regi *et al.*, 2011) to integrate DNase-seq footprints with a recalibrated position weight matrix (PWM) model for the sequence to predict the functional impact of SNPs in footprints. In CentiSNPs, SNPs in footprints “footprint-SNPs” are further categorized using CENTIPEDE hierarchical prior for each allele as “effect-SNP” if the prior relative odds for binding are *>* 20 or as “Non-effect-SNPs” otherwise.

For the other computational annotations we set the following thresholds. To run GKM-svm (Lee *et al.*, 2015), we extracted sequences around MPRA variants (19bp total) and then ran the reference vs alternate allele sequences with either the GM12878 or HepG2 weights. We then used a threshold of <−6 or > 6 for the variant scores. DeepSEA (Zhou and Troyanskaya, 2015) variant scores were identified using the website tool with a vcf file input (containing the MPRA variants). The functional significance predictions have a threshold of *<* 0.05. We overlapped SNPs from MPRA counts data with each annotation type. To identify particular annotations that predict the ASE found in the MPRA, we built logistic models log(*p*_*l*_*/*(1 *-p*_*l*_)) =*β*_0_ + *β*_1_ × *a*_*l*_ using the QuASAR *p*-values (*p <* 0.001) as the observed binary outcome, and the genomic annotations *a*_*l*_ as the predictor. For this type of analysis we use the nominal p-value (*p <* 0.001), as we test for an enrichment with respect to what would be expected from the null uniform distribution (0.1% of the tests). This nominal *p*-value corresponds to a FDR threshold of 7.2% for FDR (enrichments are not sensitive to variations of this threshold). A significant *p*-value from the annotation logistic model together with the QQ-plot are useful to evaluate which annotations work best in predicting changes in gene regulation.

## 3 RESULTS

### 3.1 Applying QuASAR-MPRA to identify ASE

We used the method proposed here, QuASAR-MPRA, to detect ASE in the MPRA data collected by Tewhey *et al.*. In MPRA, ASE is defined as the departure in the RNA reads from the DNA proportion (the input allelic ratio). Because strand orientation may affect the enhancer function of the sequences tested, each SNP was tested for ASE in the two strand orientations separately (forward/reverse). The two LCL biological replicates were combined using meta-analysis (See Methods). The number of SNPs with significant ASE (10% FDR) were 309 (forward) and 293 (reverse) in LCLs (Table S2 and Figure 1), 85 (forward) and 84 (reverse) in HepG2 (Table S3 and Figure S1). We then compared these results to those obtained using other methods previously used for MPRA/ASE analysis using the same input file with the same pre-processing filters (see Methods).

**Fig. 1.**
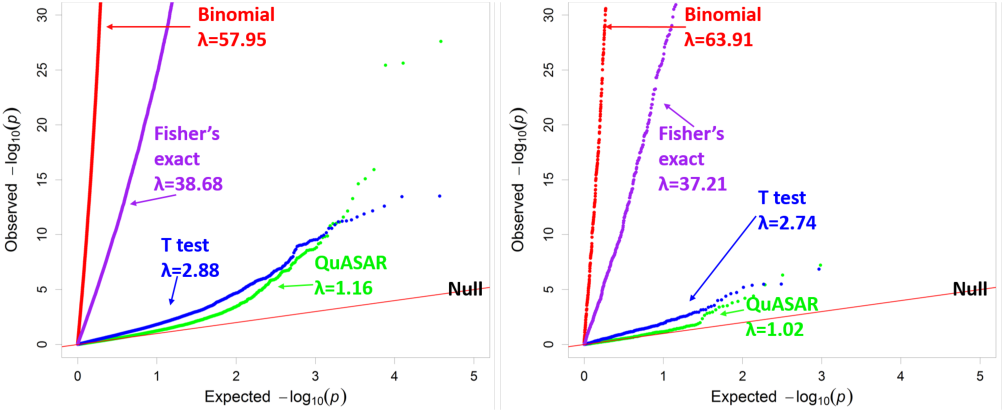
Comparing ASE testing methods in LCLs from Tewhey *et al.*. QQplot depicting the *p*-value distributions from testing for ASE using four different methods in LCLs with all SNPs (Left) or SNPs predicted to not have any regulatory effect (non-effect SNPs, Right). *λ* measures genomic inflation deviation from the uniform.

While some of the other methods seem to identify a larger number of SNPs with significant ASE, the distribution of *p*-values (Figure1) shows that those methods have very skewed distributions. The majority of genetic variants tested are expected to have no impact and only those that were the truly causal eQTL SNP should have a significant *p*-value. We do not know a priori which variants have ASE, but in Figure 1 we would expect that the majority of *p*-values would follow the expected uniform distribution if the approach correctly models the data under the null hypothesis. In other words, only a fraction of MPRA constructs are expected to have significant allelic effects. To better quantify the departure from the expected distribution of *p*-values for each testing method we used the genomic inflation method. In this method, a greater departure from a lambda value of 1 corresponds to greater inflation in the test results (see Supplement for reverse oligo results). Based on the genomic inflation value *λ*, QuASAR-MPRA results in the lowest inflation, with *λ* = 1.161. A paired *t*-test with independent estimation of variance and Welch’s adjustment, as in Tewhey *et al.*, results in a moderately but significantly larger, *λ* = 2.89 (*p <*2.2 10^−16^). The binomial test produces the greatest inflation, with *λ* = 57.95, followed by Fisher’s exact test, as in (Vockley *et al.*, 2015) resulting in *λ* = 38.68. The methods with the lowest inflation, QuASAR-MPRA and the *t*-test, have a 69% match at 10% FDR.

These results are also similar if we use a different dataset (Ulirsch *et al.*, 2016) (Figure 2). QuASAR-MPRA results in the lowest inflation, with *λ* = 0.58, while the binomial test produces the greatest inflation, with *λ* = 33.31 followed by Fisher’s exact test *λ* = 16.86. The paired *t*-test is relatively well calibrated *λ* = 1.74 but detects less hits than QuASAR-MPRA (only 64 constructs containing 53 variants at FDR 10%). Using QuASAR-MPRA we were able to identify 103 constructs containing 95 variants (FDR 10%) with significant ASE.

**Fig. 2.**
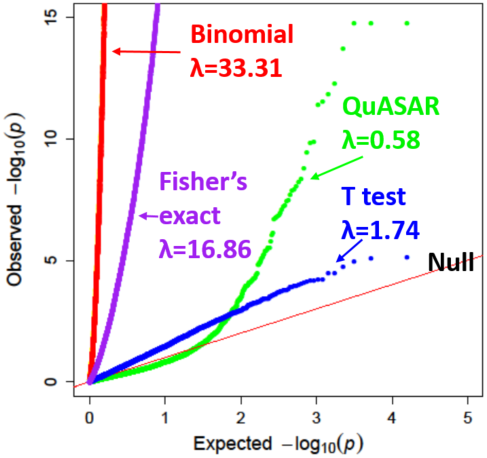
Comparing ASE testing methods in Ulirsch *et al.* dataset. QQplot depicting the *p*-value distributions from testing for ASE using four different methods in K562 for all SNPs. *λ* measures genomic inflation deviation from the uniform.

Alternatively, we also considered the *p*-value distributions only for the SNPs not predicted to affect TF binding (non-effect SNPs), as these SNPs are more likely to be true negatives 1. Note that our computationally predicted effects are not a perfect gold standard and in fact one major application of this type of data and its analysis is to precisely validate the accuracy of these computational annotations and predictions as we will show later. Nevertheless, we see (in Figure 1, 2 and S1) that the two methods with lowest lambda values show an even lower departure from the null, consistent with the computational method correctly predicting a large number of true positives.

### 3.2 Applying QuASAR-MPRA to simulated data

To further investigate our proposed new method we used simulated data where we know exactly the underlying true ASE signal to evaluate the detection accuracy. It is important to note that the simulation conditions may not exactly match those from the real data (see Methods) but they are very useful for getting more insights about the scenarios that may have larger impact on performance. Here we only compare the two methods that seem to be well-calibrated under the null hypothesis QuASAR-MPRA and the *t*-test. Under the null distribution for all our simulations both tests do not show a significant departure from the expected uniform distribution for the *p*-values.

We then compared results from QuASAR-MPRA and the *t*-test in scenarios when a fraction of the tests do have ASE (see methods). In every condition QuASAR-MPRA has greater sensitivity to detect ASE than the *t*-test (Figure 3). The *t*-test seems to perform better when the over-dispersion is low (*M* =100), or when the effect size of ASE is high (Δ*β*=2). QuASAR-MPRA also handles well low coverage data and a small number of replicates to achieve good statistical power (Figure 3). This is consistent with our original findings with QuASAR (Harvey *et al.*, 2014) demonstrating that we can measure ASE in a small number of replicates if there is enough read coverage. The *t*-test appears to require a large number of replicates in order to have power to detect ASE as compared to QuASAR-MPRA.

**Fig. 3.**
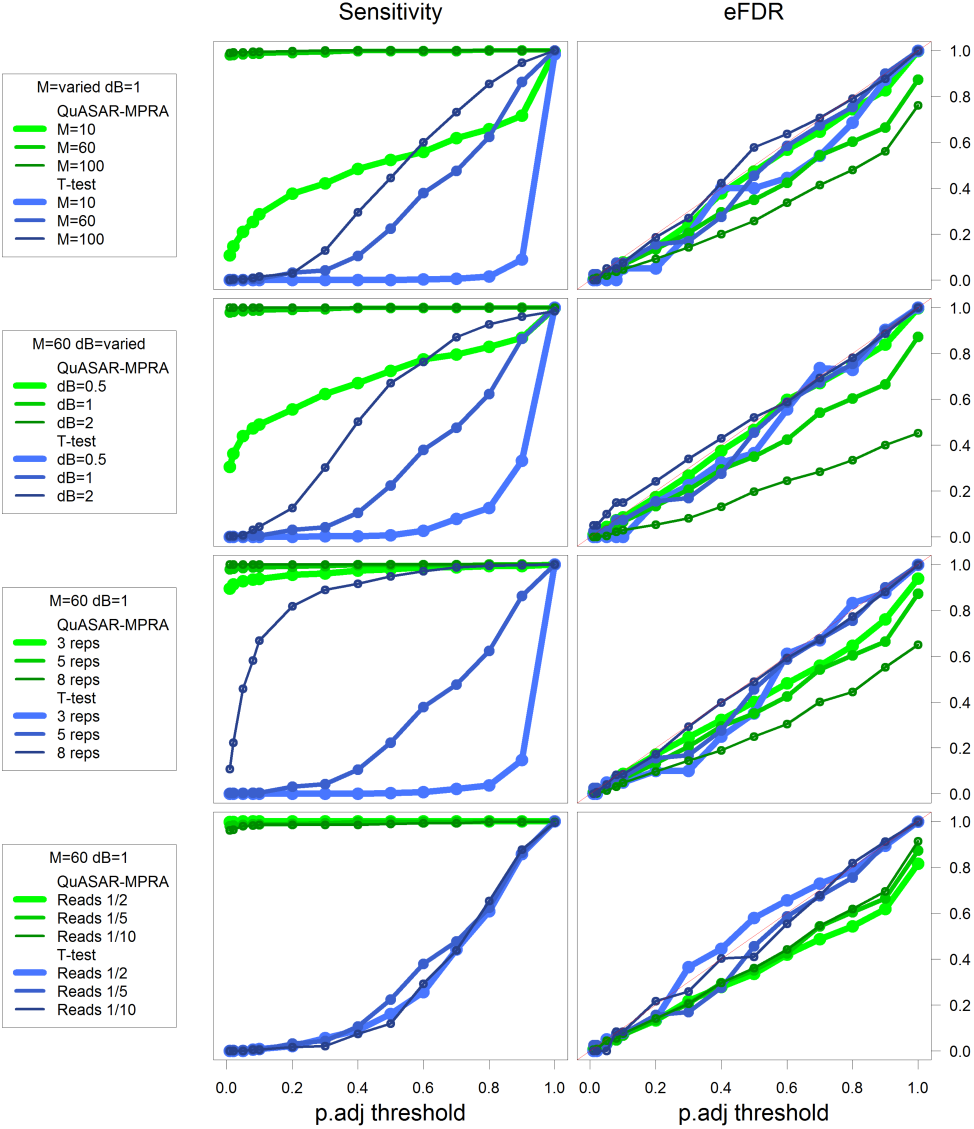
Exploring the performance across multiple simulated conditions. Plots depicting empirical power (sensitivity, Left *y*-axis) and empirical FDR (eFDR, Right *y*-axis) achieved at different BH-FDR control levels (*x*-axis) for ASE testing using QuASAR-MPRA (green) and a *t*-test (blue) across multiple simulated conditions (rows). Default conditions are M=60, Δ*β*=1, 5 replicates, and reads/5. Each row explores changing different simulation settings: A) over-dispersion high (M=10), medium (M=60) and low (M=100); B) effect-size high (Δ*β* =2), medium (Δ*β* =1) and low (Δ*β* =0.5); C) number of replicates (3, 5, or 8) D) overall sequencing depth compared to Tewhey *et al.* (1/10, 1/5, or 1/2).

### 3.3 Validation of experimental and computational annotations for functional non-coding variants

High-throughput reporter assays can be used not only to fine-map causal variants in both GWAS and eQTL studies, but also to validate SNP functional annotations (Kwasnieski *et al.*, 2014). Here we take advantage that the *p*-values derived from QuASAR are well calibrated under the null hypothesis to examine enrichments for low *p*-values in both experimentally and computationally derived annotations for allele-specific effects on TF binding. The experimentally derived annotations included LCL dsQTLs (Degner *et al.*, 2012), allele-specific hypersensitivity (ASH) SNPs (Moyerbrailean *et al.*, 2016b), and GTEx eQTLs (Consortium *et al.*, 2015). In both LCLs (Figure 4) and HepG2 (Figure S2), ASH SNPs had the greatest departure from the null, followed by LCL dsQTLs.

**Fig. 4.**
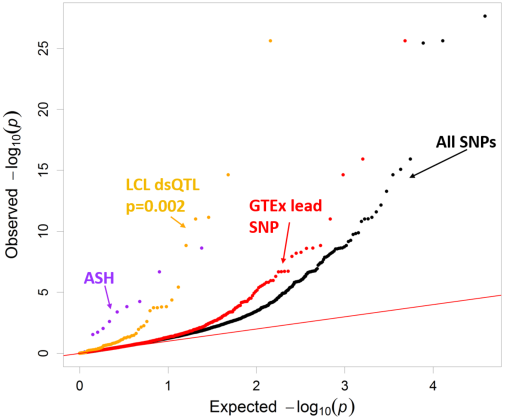
Validating experimental annotations in LCLs. QQ plot depicting the *p*-value distributions from testing for ASE using QuASAR, overlapping with experimental genomic annotations including allele-specific hypersensitivity (ASH) (Moyerbrailean *et al.*, 2016b), DNase I sensitivity QTLs (dsQTLs) (Degner *et al.*, 2012) and GTEx (Genotype-Tissue Expression) lead SNP in LCLs (Consortium *et al.*, 2015). An annotation enrichment *p*-value is reported next to their labels, but only for those annotations that are significantly enriched for small QuASAR-MPRA *p*-values according to the logistic model (see Methods).

We then asked which computational annotations seem to be the most complete and accurate predictors of the effect of a sequence variant on gene regulation as validated by MPRA. We considered effect-SNPs active in LCLs or HepG2 (Moyerbrailean *et al.*, 2016b), non-effect SNPs (negative control) (Moyerbrailean *et al.*, 2016b), predicted functional SNPs from CATO (Maurano *et al.*, 2015), GKM-svm (Lee *et al.*, 2015) (a gapped kmer sequence-based computational method to predict the effect of regulatory variation), and DeepSEA (Zhou and Troyanskaya, 2015) (predicts genomic variant effects at the variant position using deep learning-based approach). Each of the functional annotations show marked differences in *p*-value distribution. As expected, SNPs in active TF footprints, but not predicted to affect binding, show no departure from the overall distribution. In both LCLs (Figure 5) and HepG2 (Figure S3), CATO and GKM-svm SNPs had the greatest departure from the null, closely followed by effect-SNPs.

**Fig. 5.**
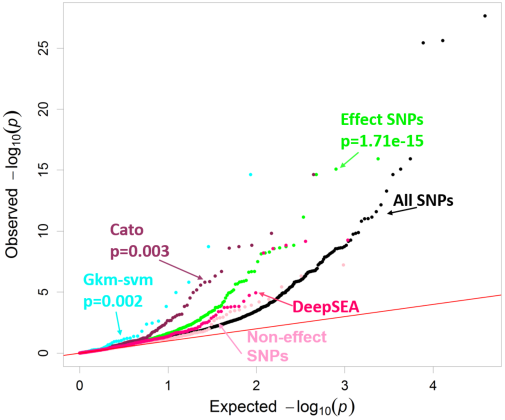
Validating computational genomic annotations in LCLs. QQ plot depicting the *p*-value distributions from testing for ASE using QuASAR, overlapping with computational genomic annotations in LCLs. Effect-SNP scores have a threshold of < −3 or *>* 3. CATO Maurano *et al* (2015) prediction scores have a threshold of *>* 0.1. GKM-svm Lee *et al.* (2015) gapped kmer sequence-based computational method to predict the effect of regulatory variation has a threshold of < −6 or > 6. DeepSEA Zhou and Troyanskaya (2015) predicts genomic variant effects at the variant position using deep learning-based algorithmic framework. The functional significance predictions have a threshold of *<* 0.05. An annotation enrichment *p*-value is reported next to their labels, but only for those annotations that are significantly enriched for small QuASAR-MPRA *p*-values according to the logistic model (see Methods).

However, effect-SNPs annotated a considerably larger number of SNPs for both cell-types and were also able to predict cell type-specific effects. LCL effect-SNPs in LCLs had a *p*-value distribution with a greater departure from the null than the HepG2 effect-SNPs (Figure S4) (*p* = 1.77 × 10^−15^ for LCL effect-SNPs vs *p*=0.14 for HepG2 effect-SNPs), whereas HepG2 effect-SNPs in HepG2 had a *p*-value distribution with a greater departure from the null than the LCL effect-SNPs (*p* = 1.81 10^−4^ for HepG2 effect-SNPs vs *p* = 1.06×10^−7^ for LCL effect-SNPs Figure S5). The differences found here in HepG2 however are minor, potentially due to fewer annotations (993 annotated LCL effect-SNPs vs 193 HepG2 effect-SNPs).

Finally, to formally quantify which annotations are the best predictors of the ASE found in the MPRA, we used all experimental and computational annotations within a logistic model to predict which SNPs in the MPRA data have a nominally significant QuASAR *p*-value (*p <* 0.001). The top predictors were GKM-svm SNPs (*p <* 2 10^−16^) and effect-SNPs (*p* = 2.17 × 10^−15^) in LCLs (Table S4). In HepG2, effect-SNPs were the greatest predictor (*p* = 1.18 × 10^−10^) (Table S5).

## 4 DISCUSSION

High throughput reporter assays have proven extremely useful for the experimental validation of enhancer regions. The recent adaptation of MPRA to investigate ASE additionally allows for validation of regulatory variants in TF binding sites, which have been shown to be functionally relevant to fine map eQTLs and GWAS signals. These large datasets, however, require analysis methods to handle the intrinsic overdispersion resulting from the original plasmid proportions, variability in the allelic imbalance, and base-calling errors.

The major advantage of QuASAR-MPRA compared to other well calibrated methods is that it requires a small number of replicates allowing for a more efficient study design. QuASAR-MPRA (along with the other methods used here) resulted in a computation time of under a minute and should scale linearly with the number of SNPs being tested. Our QuASAR-MPRA approach identifies causal regulatory variants from high-throughput reporter assays by taking into account overdispersion present in the data. This results in a well calibrated test, with minimal inflation, as determined by lambda values close to 1. In addition to being a robust method to identify ASE in high throughput reporter assays, this method estimates effect sizes and standard errors for each SNP, which can be used in fixed effects meta-analysis to easily combine datasets. Additionally, we retain a larger number of discoveries 602 (FDR 10%) compared to the original MPRA study (441 at 10%FDR) in LCLs.

Finally, we show that the allele-specific regulatory functions identified with QuASAR-MPRA can be used to validate genomic annotations as predictors for allele-specific effects. Knowing which annotations are the best predictors can aid in identifying true causal SNPs. Here we find that LCL dsQTLs and effect-SNPs are significantly predictive of ASE in LCLs and HepG2 with CATO, while GKM-svm is significant in only LCLs. Using genomic annotations can additionally help us assign mechanism of action to these regulatory variants. If a variant impacts a TF binding site for example, this can lead to gene expression changes, and therefore phenotypic effects. The less compelling results found in HepG2 may be due to HepG2 having fewer RNA replicates in the MPRA dataset than LCLs. Also there is less data available by ENCODE for the various genomic annotations, likely due to the fact that LCLs (particularly GM12878) are a tier 1 cell line and that other studies also used it for dsQTL analysis (Degner *et al.*, 2012), where HepG2 is only a tier 2 cell type.

Here we have used QuASAR-MPRA on two MPRA datasets, however this method can potentially be used for other high-throughput reporter assays, such as the ones derived from the STARR-seq protocol (e.g., POP-STARR-seq) (Vockley *et al.*, 2015) and CRE-seq protocols (Kwasnieski *et al.*, 2012), and in the context of high-throughput mutagenesis experiments. As the quest for functional validation of regulatory variants becomes more and more wide-spread, these high throughput reporter assays, when combined with a robust statistical test, represent a unique resource to functionally characterize genetic variants at an unprecedented and expandable scale.

## ACKNOWLEDGEMENT

We would like to thank Wayne State University HPC Grid for computational resources and members of the Luca/Pique group for helpful comments and discussions. Additionally we would like to thank the reviewers for their comments and suggestions.

## Funding

NIH 1R01GM109215-01 (RPR, FL)

AHA 14SDG20450118 (FL) and AHA 17PRE33460295 (CK)

